# eDNA Captures Microhabitat Partitioning in a Kelp Forest Ecosystem

**DOI:** 10.1101/2021.06.01.446542

**Authors:** Keira Monuki, Paul H. Barber, Zachary Gold

## Abstract

Environmental DNA (eDNA) metabarcoding is an increasingly important tool for surveying biodiversity in marine ecosystems. However, the scale of temporal and spatial variability in eDNA signatures, and how this variation may impact eDNA-based marine biodiversity assessments, remains uncertain. To address this question, we systematically examined variation in vertebrate eDNA signatures across depth (0 m to 10 m) and horizontal space (nearshore and surf zone) over three successive days in a Southern California kelp forest. Across a broad range of marine vertebrates (teleosts, elasmobranchs, birds, and mammals), results showed significant variation in species richness and community assemblages across 4-5 m depth, reflecting microhabitat depth preferences of common Southern California nearshore rocky reef taxa. We also found significant differences in community assemblages between nearshore and surf zone sampling stations at the same depth, and across three sampling days. Patterns of microhabitat partitioning in eDNA signatures across space and time were largely consistent with known habitat preferences and species behavior. Results highlight the sensitivity of eDNA in capturing fine-scale vertical, horizontal, and temporal variation in marine vertebrate communities, demonstrating the ability of eDNA to capture a highly localized snapshot of marine biodiversity in dynamic coastal environments.

## Introduction

Environmental DNA, or eDNA, is a metabarcoding approach for detecting and cataloging local community biodiversity, and is increasingly used in marine habitats [1,2]. eDNA approaches rely on the capture of freely dissociated cells/DNA originating from organisms in the environment, which can then be isolated and sequenced to reconstruct the community of organisms present [1,3]. Previous applications of eDNA metabarcoding in marine ecosystems demonstrate its effectiveness for measuring biodiversity and capturing important ecological information about taxa ranging from microbes to mammals [1,2], making it particularly useful for ecosystem monitoring.

The rapid rise of eDNA metabarcoding in marine ecosystems is driven by several key advantages of this technique. First, eDNA is cost-effective and has the potential for automation, allowing for increased sampling effort compared to traditional SCUBA-based underwater visual surveys [4,5]. Second, DNA barcoding approaches do not require extensive taxonomic expertise, allowing for the identification of a broad range of marine taxa from a single sample. Third, eDNA sampling only requires the collection of sea water, eliminating the potential risks associated with repeated dives needed for SCUBA-based visual surveys [4,6].

Despite these advantages, because eDNA is a relatively new technique, we know less about methodological biases compared to well-established visual survey methods. In particular, key questions remain about the temporal and spatial variability of eDNA signatures in marine environments, information that is critical for the effective design of eDNA biomonitoring efforts. Previous studies in dynamic freshwater ecosystems show daily variation in eDNA signatures [7], yet little is known about temporal variation in marine ecosystems. Degradation of eDNA signatures is relatively rapid in marine environments [8–10], with laboratory experiments showing degradation rates on the order of days [11–13]. However, in field conditions, detection of a specific eDNA point source can degrade beyond detection limits in only a few hours [14,15].

This evidence for rapid eDNA turnover rates suggests that eDNA signatures at a given location could be highly dynamic over time, particularly for species that are transitory.

In addition to temporal variability, eDNA signatures vary across horizontal space. For example, Port et al. [16] distinguished distinct communities from eDNA samples separated by only 60 m in kelp forest ecosystems, and other studies report spatial variation in eDNA signatures on similar spatial scales [17,18]. In contrast, other studies such as O’Donnell et al. [19] detect variation among communities across much larger distances, on the order of thousands of meters. A key difference in these studies is O’Donnell et al. examined eDNA from a dynamic nearshore ecosystem, whereas Port et al. examined eDNA from a highly protected cove, suggesting that physical oceanographic processes may influence the scale of eDNA variation.

Although the studies above demonstrate variation of eDNA community signatures across horizontal space, less is known about variation in marine eDNA signatures vertically across depth. In fact, most eDNA studies collect samples from consistent depths to control for depth variation [16,19,20]. The few studies that sample across multiple depths found significant variation in eDNA signatures. For example, Yamamoto et al. [17] found differences in Japanese jack mackerel eDNA concentrations between samples from 1.5 m and 31 m depth. Andruszkiewicz et al. [21] and Lacoursière-Roussel et al. [22] report similar differences in eDNA signatures between samples collected at the surface and at depth, suggesting that eDNA can resolve vertical differences in marine communities.

The scale of depth variation of eDNA signatures in these studies was multiple tens of meters, yet marine communities can partition along much smaller depth gradients in nearshore environments [23,24]. In one study, Jeunen et al. [26] found fine-scale depth variation in eDNA signatures from samples separated by only 4 vertical meters. However, these samples were taken from a fjord characterized by a strong halocline and very low wave energy, both of which greatly reduce vertical mixing. It is unclear whether similarly fine-scale vertical variation in eDNA signatures exists in exposed, nearshore coastal marine ecosystems where dynamic physical oceanographic processes should promote vertical mixing, potentially homogenizing eDNA signatures across depth [26].

To better understand the scale of spatial and temporal variation of eDNA signatures, we examined eDNA vertebrate community composition in a nearshore kelp forest and adjacent surf zone habitat in Malibu, CA where dynamic physical oceanography may homogenize eDNA community signatures. Specifically, we test 1) whether eDNA can differentiate among vertically structured vertebrate communities across a fine-scale depth gradient, 2) whether eDNA signals from similar depths vary among adjacent near-shore habitats, and 3) whether these patterns are stable over time in an exposed coastal marine environment.

## Materials and methods

We conducted our study at Leo Carrillo State Beach, Malibu, California, USA (34.0446° N, 118.9407° W). We sampled on three successive days (September 24 to September 26 in 2018) to test for temporal stability of spatial variation in eDNA signatures. On each day, we sampled at the highest tide of the mixed semidiurnal tide to minimize the impact of tidal variation on sampling.

To ensure that results reflected variation in spatial sampling, rather than time, we synchronized watches and worked in multiple teams to simultaneously sample from five depths on SCUBA along a vertical transect in a kelp forest ~140 m from shore: 0 m (at the ocean surface), 1 m, 5 m, 9 m, and 10 m (just above the sea floor). We also sampled a sixth station along the shore in the surf zone, where we collected samples approximately 1 m below the water surface where depth was approximately 2 meters. At each location, we collected triplicate seawater samples using one-liter enteral feeding bags (Kendall-Covidien – 702500) following the methods of Curd et al [27]. After returning to shore, we immediately gravity filtered all samples through 0.22 μm Sterivex filters to isolate eDNA [27]. We similarly filtered one liter of distilled water as a negative template control. Upon completion of filtration, we stored the dried filters at −20°C until extraction 48 hours later.

We extracted the DNA from the filters at UCLA using the Qiagen DNEasy Blood and Tissue kit (Qiagen, Valencia, CA, USA). To maximize eDNA recovery, we employed modifications made by Spens et al. [28], adding proteinase K and ATL buffer directly to the filter cartridges before overnight incubation in a rotating incubator at 56°C. We amplified the extracted eDNA using the *12S* MiFish Universal Teleost (MiFish-U) and MiFish Elasmobranch (MiFish-E) primers with linker modifications for Nextera indices (Table S1) [29]. Though the primers target teleost fish and elasmobranchs, they can also amplify other vertebrate species such as birds and mammals [29,30]. PCR amplification and library preparation was conducted following the methods of Curd et al. [27] (Appendix S1). After library preparation, we sequenced the library on a NextSeq at the Technology Center for Genomics & Bioinformatics (University of California, Los Angeles, CA, USA) using Reagent Kit V3 with 30% PhiX added to the sequencing run.

We processed the resulting sequencing data using the *Anacapa Toolkit* (version 1.0) for quality control, amplicon sequence variant parsing, and taxonomic assignment using standard parameters [27]. The *Anacapa Toolkit* sequence QC and ASV parsing module relies on *cutadapt* (version 1.16) [31], *FastX-toolkit* (version 0.0.13) [32], and *DADA2* (version 1.6) [33] as dependencies and the *Anacapa classifier* module relies on *Bowtie2* (version 2.3.5) [34] and a modified version of *BLCA* [35] as dependencies. We processed sequences using the default parameters and assigned taxonomy using two *CRUX-*generated reference databases. We first assigned taxonomy using the California Current Large Marine Ecosystem fish specific reference database [36]. Second, we used the *CRUX*-generated *12S* reference database supplemented with California Current Large Marine Ecosystem fish specific references to assign taxonomy using all available *12S* reference barcodes to identify any non-fish taxa following the methods of Gold et al. 2020 [35]. *CRUX* relies on *ecoPCR* (version 1.0.1) [37], *blastn* (version 2.6.0) [38], and *Entrez-qiime* (version 2.0) [39] as dependencies.

The resulting *Anacapa*-generated taxonomic tables were transferred into R for further processing(https://datadryad.org/stash/share/aMH1xTddyGgAhaWYoV3kmmmWgqCzv6Lt9YtU9s4F6NA) [36,40]. We then decontaminated the taxonomic tables using methods developed by Kelly et al. [8] and McKnight et al. [41] as implemented in Gold (2020) [42], which removes sequences from index hopping and negative controls and conducts a site occupancy model to identify true rare sequences (Appendix S2).

Following decontamination, we converted the taxonomic tables into *phyloseq* objects (version 1.30.0) in *R* [43]. We analyzed the data in two ways: first using all taxa observed and second using only a subset of frequently occurring taxa present in at least 80% of replicate samples taken across all sites. We determined the 80% threshold based on the break observed in a species occurrences graph (Figure S1). Species richness and abundance analyses were conducted using both all taxa and the most frequently occurring taxa.

We analyzed the eDNA signatures vertically across depth and horizontally across nearshore vs. surf zone habitats. We analyzed differences in eDNA across depth using only the nearshore signatures (e.g. excluding the samples collected in the adjacent surf zone). To examine species richness across depths, we conducted ANOVA and post-hoc Tukey tests using eDNA read counts (see Table S2 for eDNA read counts) [44]. We then transformed eDNA read counts to eDNA index scores, which better correlates to abundance, following the methods of Kelly et al. [45] (see Table S3 for eDNA Index scores; S1 Supplemental Methods). To analyze the importance of sampling depth on eDNA vertebrate community composition, we conducted a PERMANOVA test using the *vegan* (version 2.5-6) package in *R* [44]. The PERMANOVA was run using Bray-Curtis dissimilarity and the model eDNA_Index ~ Depth + Day + Replicate. We also ran a multivariate homogeneity of group dispersions test using the betadisper function and Bray-Curtis dissimilarity using *vegan*. We then ran a Mantel Test and non-metric multi-dimensional scaling (NMDS) using *vegan* on Bray-Curtis dissimilarities to assess community composition differences across the depth gradient. We further analyzed vertical depth community composition by generating a gradient forest model using the *gradientForest* package (version 0.1-17) using 500 runs [46] for the taxa with the highest occurrences, subsequently referred to as “core taxa”. The environmental variables in the vertical depth gradient forest model included sampling depth, sampling day, and replicate. We then extracted the taxa with the highest model performances and plotted their eDNA index values across depth.

To analyze differences across horizontal space, we ran Welch t-tests, PERMANOVA and betadisper tests, only including samples taken at 1 m depths from nearshore and surf zone habitats. We used the Bray-Curtis dissimilarity for both tests and the model eDNA_Index ~ Location + Day + Replicate for the PERMANOVA. We also ran an additional gradient forest model with the environmental variables sampling depth, nearshore vs. surf zone, sampling day, and replicate for eDNA index scores from all stations. We then extracted the top performing taxa and plotted their eDNA index distributions.

To test whether vertical and horizontal variation in eDNA signatures were consistent over time, we compared species richness across sampling days in an ANOVA framework, looking at both total community diversity as well as the eDNA index abundances for the core taxa. The linear models used for the eDNA index ANOVA tests were lm(sqrt(eDNA_Index) ~ Depth + Day + Depth:Day). eDNA Index values were transformed to meet Normality assumptions, and the subsequent p-values were adjusted for multiple comparisons using the Bonferroni correction.

## Results

We generated a total of 23,504,223 sequence reads that passed filter from the NextSeq run. After decontamination, we recovered 21,231,865 reads and 980 ASVs representing 48 families, 71 genera and 71 species. Species detected included teleost fish (n=55), elasmobranchs (n=8), marine mammals (n=2) and birds (n=6) (Table S4). Most teleost fish and elasmobranchs were demersal (rocky reef or sandy bottom species) (n=54; 86%), while the others were pelagic (n=9; 14%) (Table S4) [47]. Of the six bird species, three were seabirds (Western gull *Larus occidentalis*, Brown pelican *Pelecanus occidentalis* and Pelagic cormorant *Urile pelagicus*) and three were terrestrial birds (American golden plover *Pluvialis dominica*, Band-backed wren *Campylorhynchus zonatus*, and Ruby-crowned kinglet *Regulus calendula*). The two marine mammal species were the Bottlenose dolphin (*Tursiops truncates*) and California sea lion (*Zalophus californianus*).

A total of 22 taxa were present in at least 80% of the samples and designated as “core taxa”. These core taxa consisted of California native teleost fish, elasmobranchs and one marine mammal *(Zalophus californianus*) (Table S5). Most of the teleost fish and elasmobranchs were demersal (n=15; 71%) taxa, with the remainder being pelagic (n=6; 29%) taxa [47].

### Vertical depth comparisons

#### Species richness

When examining all taxa, species richness differed significantly across depth, with shallow sampling stations having lower species richness than deeper stations (ANOVA; p<0.001; Fig 1a). Mean species richness for the 0 m and 1 m sites were 31.4 and 32.1, respectively, while the 5 m, 9 m, and 10 m sites were 38.2, 45.7 and 44.8, respectively. Results were similar for the core taxa, although the pattern was less pronounced (ANOVA; p=0.007; Fig 1b).

**Fig 1.**
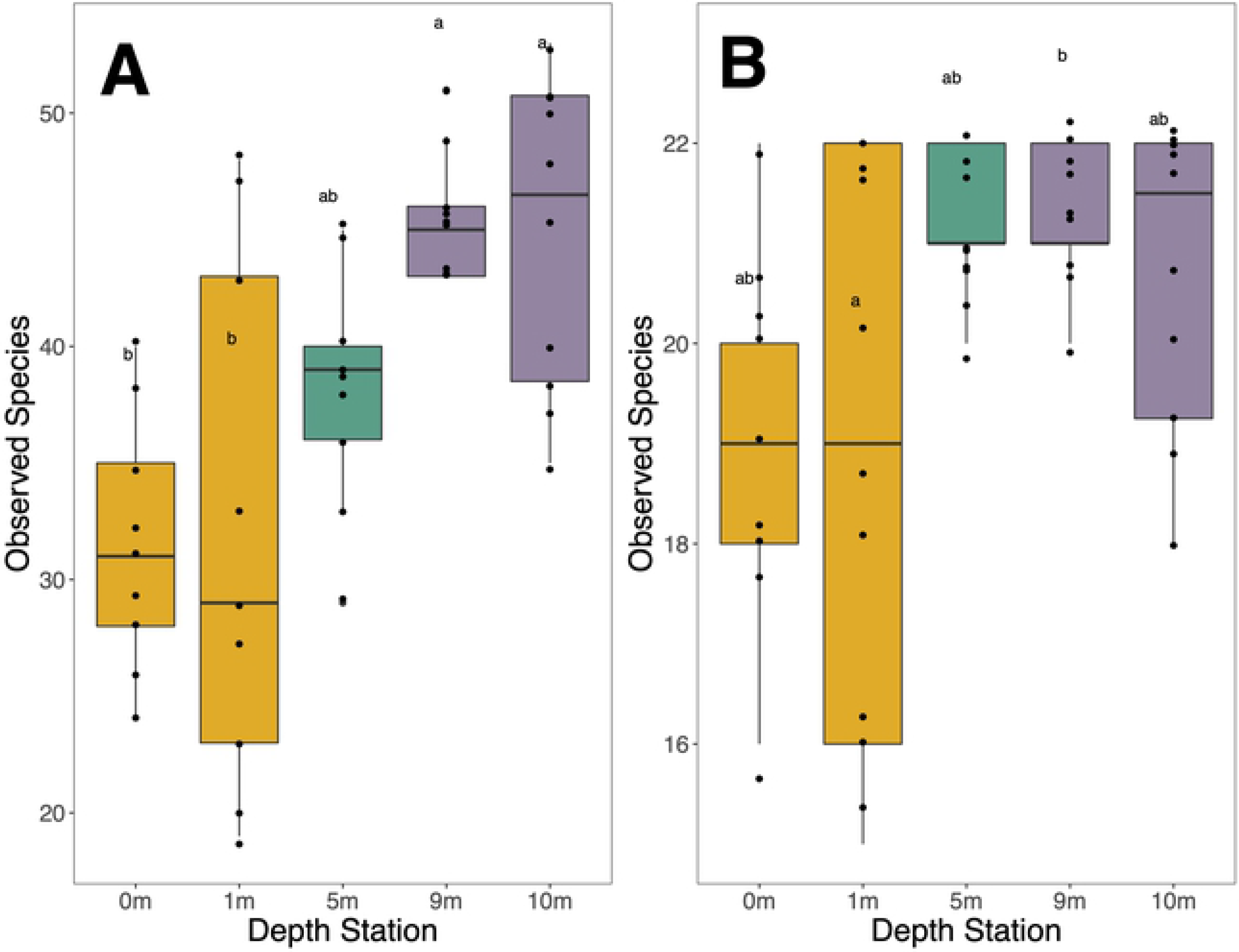
Boxplots of species richness across depth stations. The deep stations had higher species richness than the shallow stations. A) shows species richness all taxa and B) shows species richness for the high occurrence core taxa. The colors represent relative depths of the stations: yellow is shallow (0 m and 1 m), green is mid-water (5 m), and purple is deep (9 m and 10 m). See Table S6 for all taxa post-hoc Tukey test p-values and Table S7 for core taxa post-hoc Tukey test p-values.

#### Community composition

For all taxa, community composition differed significantly across depth (PERMANOVA; p=0.001). Depth accounted for most of the variation in community composition (R^2^=0.16), followed by sampling day (R^2^=0.14) and bottle replicate (R^2^=0.03) (Figure S2). The results were similar when focusing on only the core taxa (PERMANOVA; p=0.001), with depth variation accounting for the greatest variance (R^2^=0.25) in community composition followed by sampling day (R^2^=0.18) and bottle replicate (R^2^=0.02) (Figure S3).

For all taxa, pairwise PERMANOVA comparisons indicated that the 5 m, 9 m and 10 m stations were all significantly different from the 0 m and 1 m stations (all p<0.05; see Table S8 for pairwise PERMANOVA p-values). The 5 m station was significantly different from the 10 m station (p<0.05) but not the 9 m station (p=0.14). Group dispersions also differed between the 1 m and 5 m stations (p=0.02; Table S9). Pairwise PERMANOVA comparisons revealed similar patterns in the core taxa (Table S10), although group dispersions did not differ significantly between the stations (Table S11).

Mantel tests indicate that community composition significantly correlated with depth for both all taxa (Mantel statistic r=0.405, p=0.001) and core taxa (Mantel statistic r=0.457, p=0.001). Further support for this result comes from the NMDS plots, which show that communities closer in vertical distance more closely resemble each other than communities separated by greater vertical distance (Fig 2).

**Fig 2.**
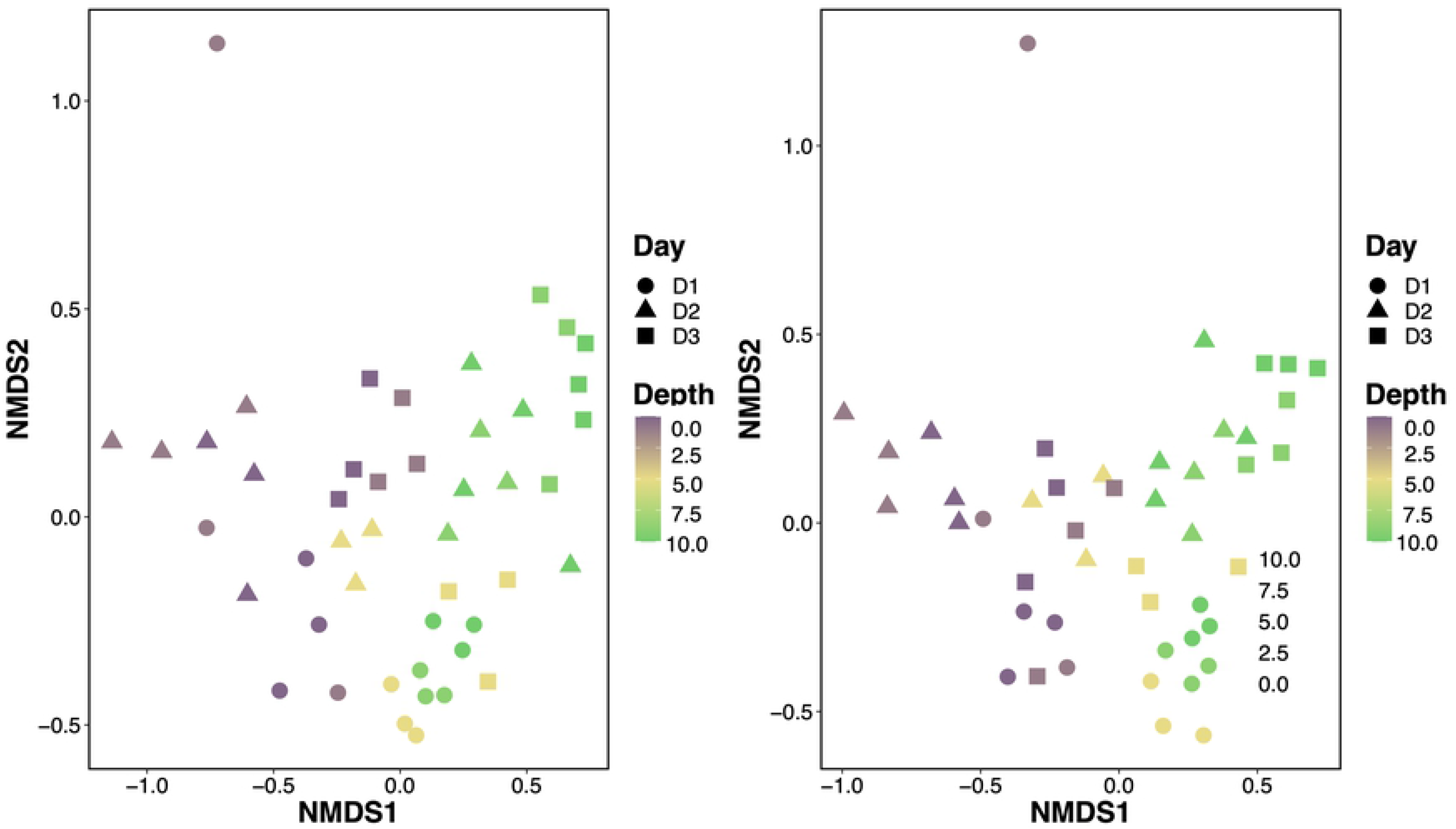
NMDS of species assemblages across depth (m) using Bray-Curtis dissimilarity. Differences in community composition increased with greater distance between depth stations for A) all taxa (stress = 0.178) and B) core taxa (stress = 0.158). 0 m represents the surface and 10 m represents the sea floor.

Of the environmental variables in the gradient forest model, depth had the highest accuracy importance (0.015) and R^2^ importance (0.183) values, followed by sampling day (accuracy importance: 0.012; R^2^ importance: 0.182) and replicate (accuracy importance: −0.003; R^2^ importance: 0.0002) (Figure S4). Of the 22 core taxa, 12 were “top predictor taxa”, characterized by high model performances (e.g. R^2^ importance > 0.34; Figure S5). Pacific sardine *Sardinops sagax,* Topsmelt silverside *Atherinops affinis* and California grunion *Leuresthes tenuis* were most abundant the shallow stations (0 m and 1 m station). The remaining nine taxa (Yellowfin drum *Umbrina roncador,* Barred sand bass *Paralabrax nebulifer,* California anchovy *Engraulis mordax,* Kelp bass *Paralabrax clathratus,* Speckled sanddab *Citharichthys stigmaeus,* California kingcroaker *Menticirrhus undulatus,* the surfperches of Family Embiotocidae, Queen croaker *Seriphus politus* and clinids of the Genus *Gibbonsia*) were most abundant in the mid and deeper stations (5 m, 9 m and 10 m) (Fig 3).

**Fig 3.**
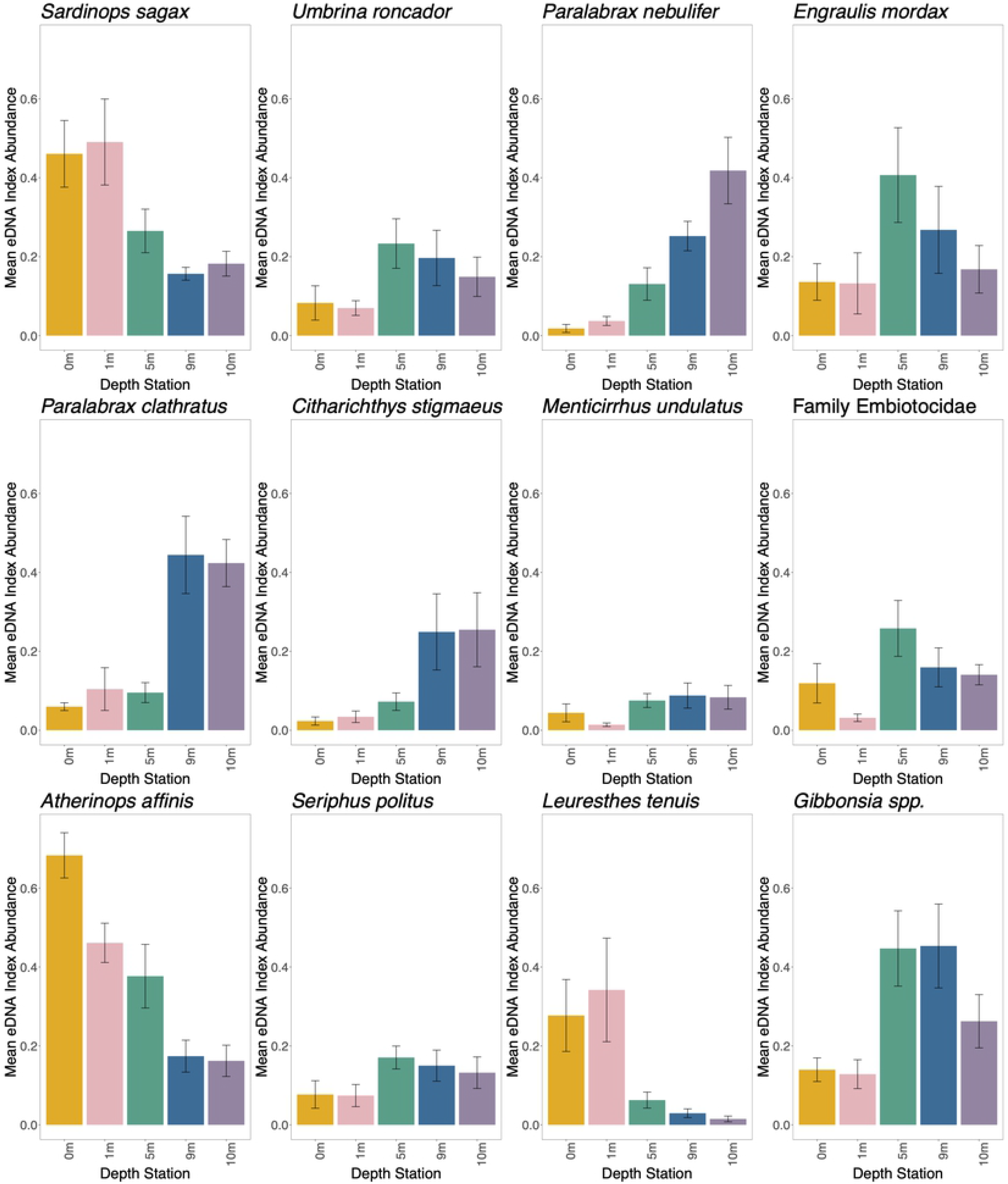
eDNA Index abundance depth distributions for the top predictor taxa. eDNA Index mean and standard error (+/− 1SE) for the top taxa (greater than 0.34 R^2^ importance) in the gradient forest model. See Supplemental Methods S2 for assigning Family Embiotocidae and Genus *Gibbonsia*.

For the remaining ten core taxa, five taxa (Kelp clingfish *Rimicola muscarum,* Kelp perch *Brachyistius frenatus,* Chub mackerel *Scomber japonicus,* Bat eagle ray *Myliobatis californica,* and California sea lion *Zalophus californianus*) were most abundant in the shallow stations and five taxa (Black perch *Embiotoca jacksoni,* Opaleye *Girella nigricans,* Giant kelpfish *Heterostichus rostratus,* Xantic sargo *Anisotremus davidsonii,* and Zebra perch sea-chub *Hermosilla azurea*) were most abundant in the mid and deep stations (Figure S6).

### Nearshore vs. surf zone analysis

#### Species richness

The surf zone station had a mean of 44.1 taxa per sample for all taxa and a mean of 19.7 taxa per sample for the core taxa compared to 32.1 and 18.9 taxa per sample, respectively, for the corresponding 1 m nearshore station. Species richness differed between the 1 m surf zone and 1 m nearshore station for all taxa (Welch two sample t-test: p=0.02) but not for core taxa (Welch two sample t-test: p=0.50).

#### Community composition

Marine vertebrate community composition differed significantly between the 1 m surf zone and 1 m nearshore stations (PERMANOVA: both all taxa and core taxa p=0.001). The 1 m surf zone station also had significantly lower group dispersions than the 1 m nearshore station for all taxa (p=0.002) and core taxa (p=0.009).

In the nearshore vs. surf zone gradient forest model, depth still had the highest accuracy importance (0.018) and R^2^ importance (0.205) values (Figure S8). Nearshore vs. surf zone designations had the second highest accuracy importance (0.011) and R^2^ importance (0.110) values, followed by sampling day (accuracy importance: 0.008; R^2^ importance: 0.081) and replicate (accuracy importance: −0.003; R^2^ importance: 0) (Figure S8). There were eight taxa with high performances in the model with > 0.40 R^2^ importance values (Figure S9). Five taxa (California kingcroaker *Menticirrhus undulatus,* surfperches of Family Embiotocidae, Yellowfin drum *Umbrina roncador,* Queen croaker *Seriphus politus* and Zebra perch sea-chub *Hermosilla azurea*) were most abundant in the surf zone station (Fig 4). Barred sand bass *Paralabrax nebulifer* and Kelp bass *Paralabrax clathratus* were most abundant in the deep nearshore stations (9m and 10 m) and Pacific sardine *Sardinops sagax* was most abundant in the shallow nearshore stations (0 m and 1 m) (Fig 4).

**Fig 4.**
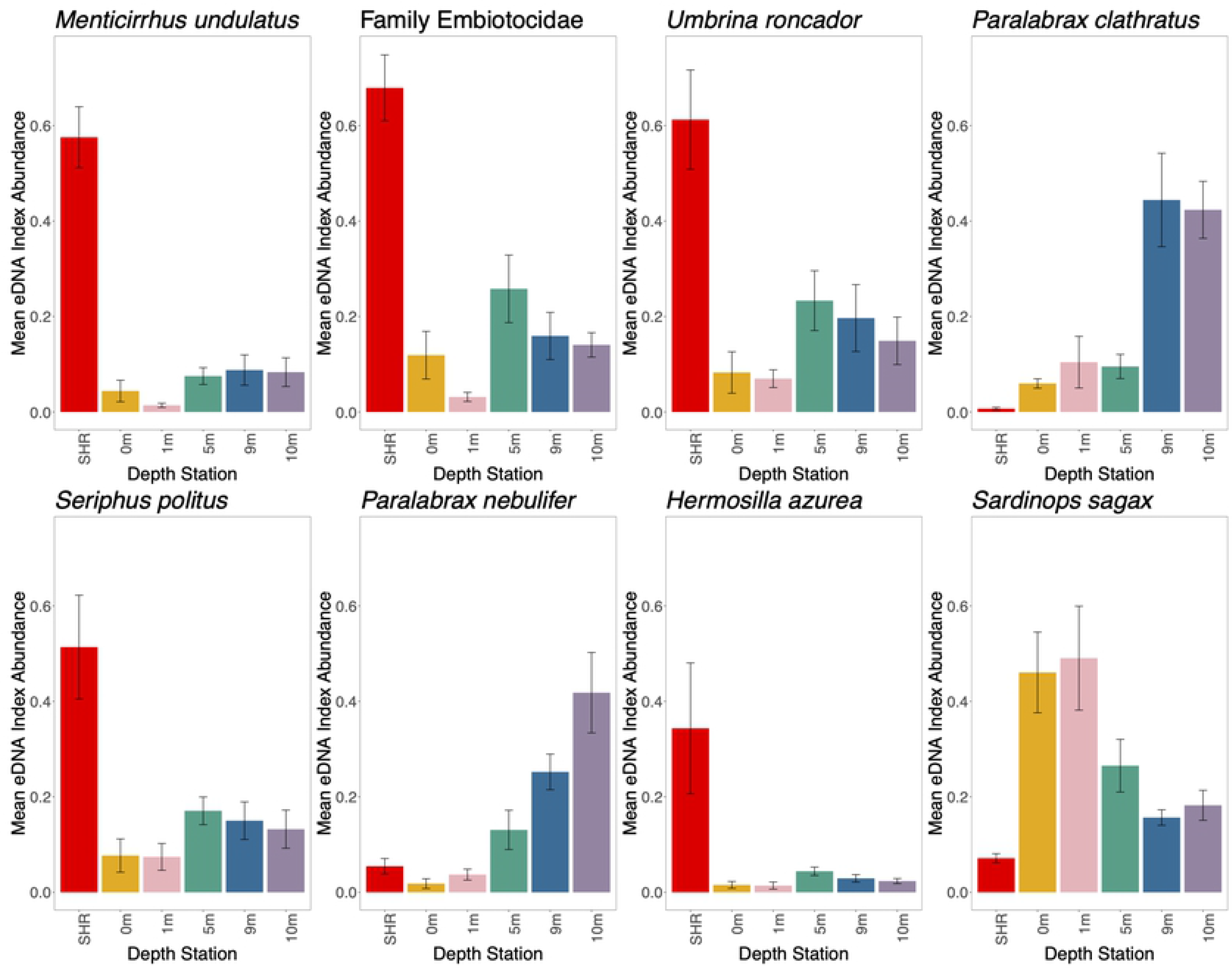
The top predictor taxa eDNA Index abundance nearshore and surf zone distributions. eDNA Index mean and standard error (+/− 1SE) for the top taxa (greater than 0.40 R^2^ importance) in nearshore vs. surf zone gradient forest model. See S2 Supplemental Methods for assigning Family Embiotocidae.

### Temporal comparisons

#### Species richness

When examining all taxa and core taxa, species richness did not differ across sampling days (all taxa ANOVA: p=0.195; core taxa ANOVA: p=0.303). All taxa mean species richness was 37.8 on Day 1, 38.1 on Day 2 and 42.6 on Day 3. Core taxa mean species richness was 19.8 on Day 1, 19.9 on Day 2 and 20.7 on Day 3.

#### Community composition

There were twelve taxa that varied strongly in the gradient model with > 0.34 R^2^ importance values (Figure S9). Six taxa (Yellowfin drum *Umbrina roncador,* California anchovy *Engraulis mordax,* California kingcroaker *Menticirrhus undulatus*, surfperches of Family Embiotocidae, Queen croaker *Seriphus politus,* and clinids of Genus *Gibbonsia*) were most abundant on Day 1, three taxa (Pacific sardine *Sardinops sagax,* Topsmelt silverside *Atherinops affinis,* California grunion *Leuresthes tenuis*) were most abundant on Day 2, and two taxa (Barred sand bass *Paralabrax nebulifer,* Speckled sanddab *Citharichthys stigmaeus*) were most abundant on Day 3 (Table S12). Kelp bass *Paralabrax clathratus* was most abundant in Day 2 and Day 3 (Table S12).

Of the 22 core taxa, 11 (50%) had depth distribution patterns that varied across the three days (all p<0.05; see Table S13 for p-values). Of these 11 taxa with variable depth distributions, six (55%) of the taxa are transitory or migratory (Queen croaker *Seriphus politus,* Yellowfin drum *Umbrina roncador,* California grunion *Leuresthes tenuis*, Pacific sardine *Sardinops sagax*, Californian anchovy *Engraulis mordax* and the surfperches of Family Embiotocidae) (Figure S10; Table S13). Of the 11 core taxa that did not have variable depth distribution patterns across the three days, seven (64%) were less mobile and less transient species (Kelp perch *Brachyistius frenatus,* Black perch *Embiotoca jacksoni*, Kelp clingfish *Rimicola muscarum,* Opaleye *Girella nigricans*, Xantic sargo *Anisotremus davidsonii*, Barred sand bass *Paralabrax nebulifer* and Speckled sanddab *Citharichthys stigmaeus*) (Figure S10: Table S13).

## Discussion

Results of eDNA surveys in a dynamic, California coastal ecosystem demonstrate fine-scale vertical and horizontal variation in marine vertebrate communities. Differences in vertebrate eDNA signatures across a 10 m depth gradient largely reflect species-specific variation in microhabitat depth preferences, particularly within fishes [48]. Similarly, eDNA discriminated between nearshore and surf zone communities with patterns also reflecting known ecological differences among fishes. While eDNA signatures varied across time, this variation was largely driven by transient taxa, while signatures of resident taxa were relatively stable. Combined, these results underscore the sensitivity of eDNA to discriminate ecologically relevant vertical, horizontal, and temporal variation.

Other studies report depth variation in eDNA over larger depth ranges [17,21], or similar scales in marine ecosystems with limited vertical mixing [25]. Studies also report horizontal partitioning on larger spatial scales along exposed coastlines [18] or on similar spatial scales in protected coves. Our results are unique in that eDNA discerned fine-scale vertical and horizontal variation in marine vertebrate communities in a dynamic, exposed coastal ecosystem, highlighting the ability of eDNA to provide a highly localized snapshot of marine diversity across depth and habitat type.

### eDNA distinguishes marine communities by depth

Previous eDNA studies report differences in eDNA signatures across depth [17,21,22,26], but these differences were observed across large depth ranges across tens of meters. The finest depth partitioning reported in an eDNA study, only 4 m [26], occurred in a unique marine ecosystem with a pronounced pyncocline driven by a 3-4 °C temperature gradient and 20-30 PSU salinity gradient [26], values that greatly exceed those observed in Southern California [49]. In such a stratified water column, in an ecosystem with limited wave energy, vertical movement of eDNA would be limited to diffusion, or potentially other processes such as biogenic vertical mixing [50,51], allowing for fine-scale variation in eDNA signatures.

While results of this study also differentiated eDNA community signatures over 4-5 m, our study site was a wave-exposed coastal environment, where currents and wave energy should facilitate vertical water mixing. Over the three sampling days of this study, daily tidal fluctuations exceeded 1 m (0.15 m low tide, 1.5 m high tide), swell height ranged from 0.23-0.43 m, and surf height ranged from 0.3-0.9 m, providing water movement and wave energy to facilitate mixing. Although we sampled at the end of the boreal summer, when the nearshore Southern California summer thermocline is typically most pronounced and within a few meters of the ocean surface [52,53], observed surface temperatures were typical for the time of year (19.2-19.4°C) and neither divers or dive computers observed a thermocline that would limit vertical mixing. As such, the community variation in eDNA signatures across a depth gradient of 4-5 m cannot be attributed to water stratification and limited mixing.

Instead, the composition of vertebrate communities detected with eDNA corresponded to species-specific depth preferences [47,48]. Of the 22 core taxa, 18 (82%) had eDNA depth distribution patterns that matched their known depth preferences (pelagic vs. demersal). For example, the pelagic species Pacific sardine *Sardinops sagax,* Topsmelt silverside *Atherinops affinis*, California grunion *Leuresthes tenuis,* Chub mackerel *Scomber japonicus* and California sea lion *Zalophus californianus* were most abundant in the 0 m and 1 m stations [48,54–56]. Thirteen demersal taxa (Barred sand bass *Paralabrax nebulifer,* clinids of the Genus *Gibbonsia*, Yellowfin drum *Umbrina roncador,* Speckled sanddab *Citharichthys stigmaeus,* California kingcroaker *Menticirrhus undulatus*, surfperches of the Family Embiotocidae, Queen croaker *Seriphus politus*, Black perch *Embiotoca jacksoni,* Giant kelpfish *Heterostichus rostratus,* and Xantic sargo *Anisotremus davidsonii,* Kelp bass *Paralabrax clathratus,* Opaleye *Girella nigricans,* and Zebra perch sea-chub *Hermosilla azurea*) were most abundant in the mid and deeper stations [48,57]. Habitat preference patterns were especially clear for the top performing species in the gradient forest model (Fig 3), indicating the species with the highest predictive power in the model are also those with eDNA distributions that most closely match their known depth partitioning. The correspondence in species-specific eDNA relative abundances and the preferred microhabitats of these species strongly suggests that eDNA is recovering ecologically informative depth variation in marine communities, adding to a growing list of studies highlighting the ability of eDNA to accurately discriminate fine-scale habitat partitioning in nearshore marine environments [48,57,58].

Species at depth greatly outnumbered species at the surface (≤1 m), with the 0 m and 1 m stations having, on average, 13 fewer taxa than the deeper stations (5 m, 9 m, 10 m) when examining all taxa. While the partitioning of shallow and deeper water vertebrate communities broadly conformed to known habitat preferences, taphonomic processes may also reduce the detection of surface-dwelling taxa. Solar radiation, particularly ultraviolet (UV) light, is highest in surface waters and may increase DNA denaturation and degradation [59–63]. Similarly, surface water temperatures are warmer, potentially increasing microbial and enzymatic activity, accelerating eDNA degradation [61,64]. Conversely, settlement processes could also elevate eDNA community diversity at depth [65]. Common sources of eDNA (e.g. feces) are often too large to remain suspended, potentially inflating eDNA community diversity at depth as these particles sink [66–68]. In our study, some pelagic, surface dwelling species were found in the deep stations, possibly due to such settlement processes. For example, Topsmelt silverside *Atherinops affinis,* California grunion *Leuresthes tenuis* and Pacific sardine *Sardinops sagax* were pelagic species found in the 9 m and 10 m stations. Though these three species were found in the deeper stations, they had low abundances in these stations and had much higher abundances in the shallow stations (Fig 3). These relative abundances reflect their habitat preferences, which suggests that, because community composition largely matches microhabitat differences, the impacts of settlement processes are likely relatively minor.

### eDNA fate and transport is likely limited

The strong depth gradient in eDNA signatures strongly suggests limited persistence and transport of eDNA, even in dynamic nearshore coastal environments. However, previous laboratory experiments and modeling studies indicate marine eDNA degradation rates on the scale of multiple days, with potential transport distances of hundreds of meters to kilometers [9,13,17,19]. One possible explanation to reconcile this apparent contradiction is high shedding rates of endogenous eDNA. If local sources of eDNA generation are high and continuous, recently generated eDNA should dominate signatures [8], as eDNA from more distant sources would have much lower concentrations due to diffusion and degradation processes during transport.

The strong eDNA signatures from a group of high occurrence core taxa observed in our study support this hypothesis. These core taxa are all abundant in nearshore rocky reef environments. Not only do the core taxa better capture eDNA variability, reducing unexplained variance from 66% in the full dataset to 55%, but they also display a more pronounced depth gradient, with depth accounting for more variation in the core taxa (25%) than all taxa (16%). Non-local, non-recent eDNA likely contributes to the high unexplained variance in our full dataset, because such eDNA may not accurately reflect the immediate environment from which it was sampled. For example, only three of the six bird species in our full dataset were seabirds; the remaining three were terrestrial bird species, whose eDNA was likely transported from external sources.

### eDNA recovers nearshore and surf zone communities

Surf zone and nearshore rocky reef community eDNA signatures varied significantly, and was comparable to variation observed by O’Donnell et al. [19] over kilometers in a dynamic coastal ecosystem. The PERMANOVA and low group dispersions for the surf zone station both indicate that the surf zone community is distinct from the nearshore community. Importantly, four of the five taxa in the nearshore vs. surf zone gradient forest model that were most abundant in the surf zone station (California kingcroaker *Menticirrhus undulatus,* surfperches of Family Embiotocidae, Yellowfin drum *Umbrina roncador,* and Queen croaker *Seriphus politus)* are all associated with surf zone habitats, providing further evidence that eDNA accurately recovers distinct communities across adjacent habitat types (Table S5) [48,57]. Combined with the above, these results indicate that eDNA can capture variation in nearshore marine ecosystems, both vertically and across habitats, highlighting that eDNA is a localized snapshot of marine diversity.

### eDNA signatures vary across time

The PERMANOVA and gradient forest analyses indicate eDNA signatures varied daily, similar to previous *in situ* eDNA studies [8–10]. Importantly, much of the temporal variance appears to result from species behavioral patterns. Of the 22 core taxa, 11 (50%) had depth distribution patterns that varied across the three days (Table S13). Of these 11 taxa, six (56%) are transitory or migratory (Table S13), including Queen croaker *Seriphus politus*, Pacific sardine *Sardinops sagax*, and the Pacific anchovy *Engraulis mordax* (Figure S10). These mobile species transit in and out of kelp forest ecosystems [48,55], potentially contributing the observed daily temporal variation in eDNA signatures. Conversely, the relative abundances of less mobile and less transient core taxa were fairly consistent over time at different depths (Figure S10). For example, the Black perch *Embiotoca jacksoni*, Kelp clingfish *Rimicola muscarum* and the Barred sand bass *Paralabrax nebulifer* had consistently higher abundances in the deep stations over the three sampling periods (Table S13).

Stability in taxa depth distributions across days may account for the high importance of depth and nearshore vs. surf zone designations in explaining eDNA variation in the gradient forest analyses. Overall, 13 of the 22 core taxa (59%) had depth distribution patterns across days that were consistent with their behavior, indicating eDNA reflects behavioral patterns of marine vertebrates within the ecosystems they inhabit (Table S13). However, the depth distributions of some of the core taxa did not match their known behavior. For example, Giant kelpfish *Heterostichus rostratus*, a non-migratory, less mobile species, had highly variable eDNA abundances across days, while Topsmelt silverside *Atherinops affinis*, a transitory species, had consistent abundances. It is unknown whether this discrepancy indicates an incomplete understanding of the ecology of these fishes, or whether physical oceanographic processes obscure potential eDNA-derived behavior patterns (e.g. algae dwelling species eDNA trapped within the boundary layer). Despite these few exceptions, the broad concordance of eDNA distributions and species ecologies underscores its ability to accurately recover marine communities over space, depth and time, opening the door to future applications of eDNA to better understand the behavior of marine organisms.

## Conclusions

Our study demonstrates the power of eDNA to distinguish unique vertebrate community microhabitats, both across depth at a single location, and across horizontally distinct communities. Patterns in spatial partitioning were relatively stable despite sampling a dynamic, nearshore marine environment, and reflected ecological differences in vertebrate communities. This consistency provides confidence for the application of eDNA methods in coastal biodiversity assessments. For example, eDNA can help us better understand species habitat distributions and how these distributions change over time with increased global change. However, our results also highlight the incredible sensitivity of eDNA metabarcoding approaches and suggest eDNA signatures only integrate biodiversity information across short time periods and small depth ranges.

These results underscore the importance of consistent sampling depth in marine eDNA studies, as variation in sampling depth could impact results. Moreover, for studies that seek to maximize sampling of local biodiversity through eDNA, efforts may need to incorporate sampling across horizontal space, depth, and time, as one depth and one location may not accurately reflect the full scope of local biodiversity of a given kelp forest. Fortunately, given the ease and cost-effective nature of eDNA sampling, such sampling efforts are not cost prohibitive, and will help us better document and monitor changes in coastal marine biodiversity.

## Acknowledgements

We thank Camille Gaynus, Nury Molina, Eric Caldera, Erick Zerecero, and Elizabeth Reid-Wainscoat for their assistance in eDNA sample collection, and thank the Barber lab undergraduates Markus Min, McKenzie Koch, Bridget Foy, Beverly Shih, Nikita Sridhar, and Cristopher Ruano for their help with laboratory work.

## Data Availability

All data is available through Dryad at https://datadryad.org/stash/share/aMH1xTddyGgAhaWYoV3kmmmWgqCzv6Lt9YtU9s4F6NA. Code used for analyses is available on GitHub at https://github.com/ksmonuki/eDNA-analysis. Raw sequences will be available on GenBank after manuscript acceptance.

## Supporting information

**Table S1. MiFish-U and MiFish-E primer sequences.**

**Table S2. Taxonomy table with read counts after decontamination.**

**Table S3. Taxonomy table with eDNA Index values after decontamination. Table S4. All taxa species list.**

**Table S5. Core taxa species list.**

**Table S6. ANOVA and post-hoc Tukey test p-values across depth stations for all taxa. Table S7. ANOVA and post-hoc Tukey test p-values across depth for the core taxa.**

**Table S8. Pairwise PERMANOVA p-values across depth for all taxa. Table S9. Group dispersions p-values across depth for all taxa.**

**Table S10. Pairwise PERMANOVA p-values across depth for the core taxa. Table S11. Group dispersions p-values across depth for the core taxa.**

**Table S12. Mean eDNA Index across days for top predictor species in the core taxa gradient forest model.**

**Table S13. Core taxa ecology, behavior and depth distribution variability across days. Table includes p-values for the Day:Depth interaction of the ANOVA for the linear model of eDNA index abundance.**

**Fig S1. Species vs. percent presence across samples.** There are 22 taxa with a percent presence ≥ 80%.

**Fig S2. Apportioned variance plot of the three variables in the PERMANOVA model for all taxa.** Depth accounted for 16% variance (p=0.001), collected date accounted for 14% variance (p=0.001) and replicate accounted for 3% variance (p=0.426).

**Fig S3. Apportioned variance plot of the three variables in the PERMANOVA model for core taxa.** Depth station accounted for the greatest variance in eDNA community composition. Depth accounted for 25% variance (p=0.001), collection date accounted for 18% variance (p=0.001) and replicate accounted for 2% variance (p=0.706).

**Fig S4. Importance of variables in the depth gradient forest model.** Sampling depth had the highest accuracy importance and R^2^ weighted importance in the core taxa vertical depth gradient model.

**Fig S5. Performance of species in the depth gradient forest model.** There were twelve top predictor species with R^2^ values greater than 0.34 in the core taxa vertical depth gradient forest model.

**Fig S6. Depth distributions of core taxa not included in the top predictor taxa in the depth gradient forest model.** There were ten core taxa not included in the top predictor taxa.

**Fig S7. NMDS for shore vs. 1 m stations.** Surf zone community composition differs greatly from the nearshore community composition. NMDS stress is 0.082.

**Fig S8. Importance of variables in the nearshore vs. surf zone gradient forest model.** Sampling depth had the highest accuracy importance and R^2^ weighted importance in the core taxa nearshore vs. surf zone space gradient forest model.

**Fig S9. Performance of species in the nearshore vs. surf zone space gradient forest model.** There were eight top predictor species with R^2^ values greater than 0.40 in the core taxa nearshore vs. surf zone gradient forest model.

**Fig S10. Depth distributions across the three sampling days.** eDNA Index abundance mean and standard error for the core taxa.

**Supplemental Methods S1.**

**Supplemental Methods S2.**

**Supplemental Methods S3.**

**Appendix S1.**

**Appendix S2.**

## References

1. Bohmann K, Evans A, Gilbert MTP, Carvalho GR, Creer S, Knapp M, et al. Environmental DNA for wildlife biology and biodiversity monitoring. Trends in Ecology & Evolution. 2014;29: 358–367. doi:10.1016/j.tree.2014.04.003

2. Stat M, Huggett MJ, Bernasconi R, DiBattista JD, Berry TE, Newman SJ, et al. Ecosystem biomonitoring with eDNA: metabarcoding across the tree of life in a tropical marine environment. Scientific Reports. 2017;7: 12240. doi:10.1038/s41598-017-12501-5

3. Taberlet P, Coissac E, Hajibabaei M, Rieseberg LH. Environmental DNA. Molecular Ecology. 2012;21: 1789–1793. doi:10.1111/j.1365-294X.2012.05542.x

4. Kelly RP, Port JA, Yamahara KM, Martone RG, Lowell N, Thomsen PF, et al. Harnessing DNA to improve environmental management. Science. 2014;344: 1455–1456. doi:10.1126/science.1251156

5. Ji Y, Ashton L, Pedley SM, Edwards DP, Tang Y, Nakamura A, et al. Reliable, verifiable and efficient monitoring of biodiversity via metabarcoding. Ecology Letters. 2013;16: 1245–1257. doi:10.1111/ele.12162

6. Denoble PJ, Marroni A, Vann RD. Annual Fatality Rates and Associated Risk Factors for Recreational Scuba Diving. Divers Alert Network; 2011. pp. 73–85.

7. Tillotson MD, Kelly RP, Duda JJ, Hoy M, Kralj J, Quinn TP. Concentrations of environmental DNA (eDNA) reflect spawning salmon abundance at fine spatial and temporal scales. Biological Conservation. 2018;220: 1–11. doi:10.1016/j.biocon.2018.01.030

8. Kelly RP, Gallego R, Jacobs-Palmer E. The effect of tides on nearshore environmental DNA. PeerJ. 2018;6:e4521. doi:DOI 10.7717/peerj.4521

9. Thomsen PF, Kielgast J, Iversen LL, Møller PR, Rasmussen M, Willerslev E. Detection of a Diverse Marine Fish Fauna Using Environmental DNA from Seawater Samples. PLOS ONE. 2012;7: e41732. doi:10.1371/journal.pone.0041732

10. Maruyama A, Nakamura K, Yamanaka H, Kondoh M, Minamoto T. Correction: The Release Rate of Environmental DNA from Juvenile and Adult Fish. PLOS ONE. 2019;14: e0212145. doi:10.1371/journal.pone.0212145

11. Sassoubre LM, Yamahara KM, Gardner LD, Block BA, Boehm AB. Quantification of Environmental DNA (eDNA) Shedding and Decay Rates for Three Marine Fish. Environ Sci Technol. 2016;50: 10456–10464. doi:10.1021/acs.est.6b03114

12. Thomsen PF, Kielgast J, Iversen LL, Wiuf C, Rasmussen M, Gilbert MTP, et al. Monitoring endangered freshwater biodiversity using environmental DNA. Molecular Ecology. 2012;21: 2565–2573. doi:10.1111/j.1365-294X.2011.05418.x

13. Collins RA, Wangensteen OS, O’Gorman EJ, Mariani S, Sims DW, Genner MJ. Persistence of environmental DNA in marine systems. Communications Biology. 2018;1: 1–11. doi:10.1038/s42003-018-0192-6

14. Murakami H, Yoon S, Kasai A, Minamoto T, Yamamoto S, Sakata MK, et al. Dispersion and degradation of environmental DNA from caged fish in a marine environment. Fish Sci. 2019;85: 327–337. doi:10.1007/s12562-018-1282-6

15. Ely T, Barber PH, Gold Z. Investigating temporal and spatial variation of eDNA in a nearshore rocky reef environment. bioRxiv. 2020; 2020.12.29.424660. doi:10.1101/2020.12.29.424660

16. Port JA, O’Donnell JL, Romero‐Maraccini OC, Leary PR, Litvin SY, Nickols KJ, et al. Assessing vertebrate biodiversity in a kelp forest ecosystem using environmental DNA. Molecular Ecology. 2016;25: 527–541. doi:10.1111/mec.13481

17. Yamamoto S, Minami K, Fukaya K, Takahashi K, Sawada H, Murakami H, et al. Environmental DNA as a ‘Snapshot’ of Fish Distribution: A Case Study of Japanese Jack Mackerel in Maizuru Bay, Sea of Japan. PLOS ONE. 2016;11: e0149786. doi:10.1371/journal.pone.0149786

18. Gold ZJ. Design and Implementation of Environmental DNA Metabarcoding Methods for Monitoring the Southern California Marine Protected Area Network. UCLA. 2020. Available: https://escholarship.org/uc/item/7zf5g0cb

19. O’Donnell JL, Kelly RP, Shelton AO, Samhouri JF, Lowell NC, Williams GD. Spatial distribution of environmental DNA in a nearshore marine habitat. PeerJ. 2017;5: e3044. doi:10.7717/peerj.3044

20. Sigsgaard EE, Nielsen IB, Bach SS, Lorenzen ED, Robinson DP, Knudsen SW, et al. Population characteristics of a large whale shark aggregation inferred from seawater environmental DNA. Nature Ecology & Evolution. 2017;1: 0004. doi:10.1038/s41559-016-0004

21. Andruszkiewicz EA, Starks HA, Chavez FP, Sassoubre LM, Block BA, Boehm AB. Biomonitoring of marine vertebrates in Monterey Bay using eDNA metabarcoding. PLOS ONE. 2017;12: e0176343. doi:10.1371/journal.pone.0176343

22. Lacoursière‐Roussel A, Howland K, Normandeau E, Grey EK, Archambault P, Deiner K, et al. eDNA metabarcoding as a new surveillance approach for coastal Arctic biodiversity. Ecology and Evolution. 2018;8: 7763–7777. doi:10.1002/ece3.4213

23. Barnes RSK, Mann KH. Fundamentals of Aquatic Ecology. John Wiley & Sons; 2009.

24. Chappuis E, Terradas M, Cefalì ME, Mariani S, Ballesteros E. Vertical zonation is the main distribution pattern of littoral assemblages on rocky shores at a regional scale. Estuarine, Coastal and Shelf Science. 2014;147: 113–122. doi:10.1016/j.ecss.2014.05.031

25. Barnes MA, Turner CR, Jerde CL, Renshaw MA, Chadderton WL, Lodge DM. Environmental Conditions Influence eDNA Persistence in Aquatic Systems. Environ Sci Technol. 2014;48: 1819–1827. doi:10.1021/es404734p

26. Jeunen G-J, Lamare MD, Knapp M, Spencer HG, Taylor HR, Stat M, et al. Water stratification in the marine biome restricts vertical environmental DNA (eDNA) signal dispersal. Environmental DNA. 2020;2: 99–111. doi:10.1002/edn3.49

27. Curd EE, Gold Z, Kandlikar G, Gomer J, Ogden M, O’Connell T, et al. Anacapa Toolkit: an environmental DNA toolkit for processing multilocus metabarcode datasets. Methods in Ecology and Evolution. 2019;Accepted Author Manuscript. doi:doi:10.1111/2041-210X.13214

28. Spens J, Evans AR, Halfmaerten D, Knudsen SW, Sengupta ME, Mak SST, et al. Comparison of capture and storage methods for aqueous macrobial eDNA using an optimized extraction protocol: advantage of enclosed filter. Methods in Ecology and Evolution. 2017;8: 635–645. doi:doi: 10.1111/2041-210X.12683

29. Miya M, Sato Y, Fukunaga T, Sado T, Poulsen JY, Sato K, et al. MiFish, a set of universal PCR primers for metabarcoding environmental DNA from fishes: detection of more than 230 subtropical marine species. Royal Society Open Science. 2015;2: 150088. doi:10.1098/rsos.150088

30. Closek CJ, Santora JA, Starks HA, Schroeder ID, Andruszkiewicz EA, Sakuma KM, et al. Marine Vertebrate Biodiversity and Distribution Within the Central California Current Using Environmental DNA (eDNA) Metabarcoding and Ecosystem Surveys. Front Mar Sci. 2019;6. doi:10.3389/fmars.2019.00732

31. Martin M. Cutadapt removes adapter sequences from high-throughput sequencing reads. EMBnet.journal. 2011;17: 10–12. doi:10.14806/ej.17.1.200

32. Gordon A, Hannon G. FASTX-Toolkit. FASTQ/A short-reads preprocessing tools (unpublished). 2010 [cited 30 Nov 2020]. Available: http://hannonlab.cshl.edu/fastx_toolkit/index.html

33. Callahan BJ, McMurdie PJ, Rosen MJ, Han AW, Johnson AJA, Holmes SP. DADA2: High-resolution sample inference from Illumina amplicon data. Nature Methods. 2016;13: 581–583. doi:10.1038/nmeth.3869

34. Langmead B, Salzberg SL. Fast gapped-read alignment with Bowtie 2. Nat Methods. 2012;9: 357–359. doi:10.1038/nmeth.1923

35. Gao X, Lin H, Revanna K, Dong Q. A Bayesian taxonomic classification method for 16S rRNA gene sequences with improved species-level accuracy. BMC Bioinformatics. 2017;18: 247. doi:10.1186/s12859-017-1670-4

36. Gold Z, Choi E, Kacev D, Frable B, Burton R, Goodwin K, et al. FishCARD: Fish 12S California Current Specific Reference Database for Enhanced Metabarcoding Efforts. Preprints; 2020 Jun. doi:10.22541/au.159136805.55528691

37. Ficetola GF, Coissac E, Zundel S, Riaz T, Shehzad W, Bessière J, et al. An In silico approach for the evaluation of DNA barcodes. BMC Genomics. 2010;11: 434. doi:10.1186/1471-2164-11-434

38. Camacho C, Coulouris G, Avagyan V, Ma N, Papadopoulos J, Bealer K, et al. BLAST+: architecture and applications. BMC Bioinformatics. 2009;10: 421. doi:10.1186/1471-2105-10-421

39. Baker C. entrez_qiime: a utility for generating QIIME input files from the NCBI databases. In: entrez_qiime [Internet]. 2016 [cited 30 Nov 2020]. Available: https://bakerccm.github.io/entrez_qiime/

40. R Core Team. R: A language and environment for statistical computing. Vienna, Austria; 2019. Available: https://www.R-project.org/

41. McKnight DT, Huerlimann R, Bower DS, Schwarzkopf L, Alford RA, Zenger KR. microDecon: A highly accurate read-subtraction tool for the post-sequencing removal of contamination in metabarcoding studies. Environmental DNA. 2019;1: 14–25. doi:10.1002/edn3.11

42. Zack Gold. zjgold/gruinard_decon: Gruinard Decon Version 0.0. Zenodo; 2020. doi:10.5281/zenodo.4291158

43. McMurdie PJ, Holmes S. phyloseq: An R Package for Reproducible Interactive Analysis and Graphics of Microbiome Census Data. PLOS ONE. 2013;8: e61217. doi:10.1371/journal.pone.0061217

44. Oksanen J, Blanchet FG, Friendly M, Kindt R, Legendre P, McGlinn D, et al. vegan: Community Ecology Package. 2019. Available: https://CRAN.R-project.org/package=vegan

45. Kelly RP, Shelton AO, Gallego R. Understanding PCR Processes to Draw Meaningful Conclusions from Environmental DNA Studies. Scientific Reports. 2019;9: 1–14. doi:10.1038/s41598-019-48546-x

46. Ellis N, Smith SJ, Pitcher CR. Gradient forests: calculating importance gradients on physical predictors. Ecology. 2012;93: 156–168. doi:10.1890/11-0252.1

47. Froese R, Pauly D. FishBase. World Wide Web electronic publication.; 2019. Available: http://www.fishbase.org

48. Eschmeyer WN, Herald ES. A Field Guide to Pacific Coast Fishes of North America. New York, NY: Houghton Mifflin Company; 1983.

49. Tapia FJ, DiBacco C, Jarrett J, Pineda J. Vertical distribution of barnacle larvae at a fixed nearshore station in southern California: Stage-specific and diel patterns. Estuarine, Coastal and Shelf Science. 2010;86: 265–270. doi:10.1016/j.ecss.2009.11.003

50. Katija K, Dabiri JO. A viscosity-enhanced mechanism for biogenic ocean mixing. Nature. 2009;460: 624–626. doi:10.1038/nature08207

51. Katija K. Biogenic inputs to ocean mixing. Journal of Experimental Biology. 2012;215: 1040–1049. doi:10.1242/jeb.059279

52. Cairns JL, Nelson KW. A description of the seasonal thermocline cycle in shallow coastal water. Journal of Geophysical Research (1896-1977). 1970;75: 1127–1131. doi:10.1029/JC075i006p01127

53. Petersen JH, Jahn AE, Lavenberg RJ, McGowen GE, Grove RS. Physical-chemical characteristics and zooplankton biomass on the continental shelf off Southern California. California Cooperative Oceanic Fisheries Investigations. 1986;27: 36–52.

54. Whitehead PJP. Vol.7. Clupeoid fishes of the world (Suborder Clupeoidei). An annotated and illustrated catalogue of the herrings, sardines, pilchards, sprats, shads, anchovies and wolf-herrings. Rome; 1985.

55. Hernández-Camacho C, Aurioles-Gamboa D. IUCN Red List of Threatened Species: Zalophus californianus. In: IUCN Red List of Threatened Species [Internet]. 12 Dec 2014 [cited 7 Feb 2021]. Available: https://www.iucnredlist.org/en

56. Collette B, Nauen C. FAO species catalogue. Vol.2. Scombrids of the world. An annotated and illustrated catalogue of tunas, mackerels, bonitos and related species known to date. [cited 19 Feb 2021]. Available: http://www.fao.org/3/ac478e/ac478e00.htm

57. Fischer W, Krupp F, Schneider W, Sommer C, Carpenter K, Niem V. Guía FAO para la identificación de especies para los fines de la pesca, Pacífico centro-oriental; vol. 3. 1995.

58. Miller DJ. Miller and Lea’s Guide to the Coastal Marine Fishes of California, 2nd Edition. UCANR Publications; 2020.

59. Kreader CA. Persistence of PCR-Detectable Bacteroides distasonis from Human Feces in River Water. Appl Environ Microbiol. 1998;64: 4103–4105.

60. Okabe S, Shimazu Y. Persistence of host-specific Bacteroides–Prevotella 16S rRNA genetic markers in environmental waters: effects of temperature and salinity. Appl Microbiol Biotechnol. 2007;76: 935–944. doi:10.1007/s00253-007-1048-z

61. Poté J, Ackermann R, Wildi W. Plant leaf mass loss and DNA release in freshwater sediments. Ecotoxicology and Environmental Safety. 2009;72: 1378–1383. doi:10.1016/j.ecoenv.2009.04.010

62. Dick LK, Stelzer EA, Bertke EE, Fong DL, Stoeckel DM. Relative Decay of Bacteroidales Microbial Source Tracking Markers and Cultivated Escherichia coli in Freshwater Microcosms. Appl Environ Microbiol. 2010;76: 3255–3262. doi:10.1128/AEM.02636-09

63. Ravanat J-L, Douki T, Cadet J. Direct and indirect effects of UV radiation on DNA and its components. Journal of Photochemistry and Photobiology B: Biology. 2001;63: 88–102. doi:10.1016/S1011-1344(01)00206-8

64. Corinaldesi C, Beolchini F, Dell’anno A. Damage and degradation rates of extracellular DNA in marine sediments: implications for the preservation of gene sequences. Molecular Ecology. 2008;17: 3939–3951. doi:10.1111/j.1365-294X.2008.03880.x

65. Stewart KA. Understanding the effects of biotic and abiotic factors on sources of aquatic environmental DNA. Biodivers Conserv. 2019;28: 983–1001. doi:10.1007/s10531-019-01709-8

66. Turner CR, Uy KL, Everhart RC. Fish environmental DNA is more concentrated in aquatic sediments than surface water. Biological Conservation. 2015;183: 93–102. doi:10.1016/j.biocon.2014.11.017

67. Turner CR, Barnes MA, Xu CCY, Jones SE, Jerde CL, Lodge DM. Particle size distribution and optimal capture of aqueous macrobial eDNA. Methods in Ecology and Evolution. 2014;5: 676–684. doi:10.1111/2041-210X.12206

68. Klymus KE, Richter CA, Chapman DC, Paukert C. Quantification of eDNA shedding rates from invasive bighead carp Hypophthalmichthys nobilis and silver carp Hypophthalmichthys molitrix. Biological Conservation. 2015;183: 77–84. doi:10.1016/j.biocon.2014.11.020

